# SdhA interferes with disruption of the *Legionella*-containing vacuole by hijacking the OCRL phosphatase

**DOI:** 10.1101/2020.10.04.325654

**Authors:** Won Young Choi, Elizabeth A. Creasey, Martin Lowe, Ralph R. Isberg

## Abstract

*Legionella pneumophila* grows intracellularly within a replication vacuole via action of Icm/Dot-secreted proteins. One such protein, SdhA, maintains the integrity of the vacuolar membrane, thereby preventing cytoplasmic degradation of bacteria. We show here that SdhA binds and blocks the action of OCRL (OculoCerebroRenal syndrome of Lowe), an inositol 5-phosphatase pivotal for controlling endosomal dynamics. OCRL depletion resulted in enhanced vacuole integrity and intracellular growth of a *sdhA* mutant, consistent with OCRL participating in vacuole disruption. Overexpressed SdhA altered OCRL function, enlarging endosomes, driving endosomal accumulation of PI(4,5)P_2_, and interfering with endosomal trafficking. SdhA interrupted Rab GTPase-OCRL interactions by binding to the OCRL ASH domain, without directly altering OCRL 5-phosphatase activity. The *Legionella* vacuole encompassing the *sdhA* mutant accumulated OCRL and endosomal antigen EEA1, consistent with SdhA blocking accumulation of OCRL-containing endosomal vesicles. Therefore, SdhA hijacking of OCRL is associated with blocking trafficking events that disrupt the pathogen vacuole.

## Introduction

*Legionella pneumophila* is the causative agent of the potentially fatal Legionnaire’s disease, growing within alveolar macrophages as a central step in its pathogenesis (Copenhaver et al., 2014; Nash et al., 1984). As an environmental bacterium, the primary selective force for intracellular growth is its ability to infect amoebae, which can contaminate a variety of plumbing and cooling systems that act as disease reservoirs (Muder et al., 1986; Rowbotham, 1980). Human infection occurs by accidental inhalation or aspiration of contaminated aerosolized water followed by intracellular growth of *Legionella* in alveolar macrophages (Horwitz and Silverstein, 1980).

The intracellular growth of *L. pneumophila* depends on the construction of the *Legionella-*containing vacuole (LCV). Once internalized, the bacterium translocates about 300 bacterial proteins into the host via the Icm/Dot Type IV secretion system (T4SS) (Huang et al., 2011; Luo and Isberg, 2004; Zhu et al., 2011). The secretion of bacterial effector proteins into the host cell allows hijacking of host membrane trafficking pathways to remodel the LCV into a membranous compartment that supports intracellular replication (Berger and Isberg, 1993; Segal, 2013; Segal and Shuman, 1999). In contrast to phagocytic uptake of nonpathogens, which is characterized by interactions with the endocytic pathway and subsequent targeting to lysosomal compartments, the LCV recruits components of the early secretory pathway, allowing direct interaction with the endoplasmic reticulum (ER) (Clemens et al., 2000; Kagan and Roy, 2002; Swanson and Isberg, 1995; Tilney et al., 2001). This ER-encompassed compartment, protected from lysosomal degradation, also sequesters the bacterium from the cytoplasmic innate immune sensing system in mammalian hosts. The extreme restriction of bacteria that enter the mammalian cell cytosol was first demonstrated by the behavior of *L. pneumophila sdhA* mutants, which have disrupted vacuoles that result in bacterial exposure to the host cytosol (Aachoui et al., 2013; Creasey and Isberg, 2012; Ge et al., 2012).

The SdhA protein is a T4SS substrate essential for intracellular growth of *L. pneumophila* in primary macrophages (Laguna et al., 2006). Release of bacteria into the mammalian cytosol in the absence of SdhA occurs via an unknown pathway, and results in recognition by cytosol-localized interferon (IFN)-stimulated anti-microbial GBPs (Guanylate Binding Proteins) leading to bacterial degradation (Liu et al., 2018; Pilla et al., 2014). The degraded bacteria release bacterial components such as LPS and DNA, which in turn activate AIM2, caspase-11, and caspase-1 inflammasomes causing pyroptotic death of the infected host cells (Creasey and Isberg, 2012; Ge et al., 2012; Pilla et al., 2014). Therefore, even if the vacuole avoids entry into the lysosomal pathway, disruption of the vacuole can lead to cytosolic bacterial degradation. RNAi depletion of Rab5, Rab11, and Rab8, all GTPases involved in endocytic and recycling pathways, partially reverses loss of vacuole integrity observed in *sdhA* mutants. Consistent with these results, the absence of SdhA results in LCV accumulation of EEA1 and Rab11FIP1, downstream effectors of these GTPases (Anand et al., 2020; Christoforidis et al., 1999; Hales et al., 2001). Therefore, it is likely that SdhA interferes with components of the early endocytic network that are likely to disrupt vacuole integrity.

One protein involved in controlling the identities of compartments associated with the endocytic network is OCRL (OculoCerebroRenal syndrome of Lowe), a polyphosphoinositide-5-phosphatase that regulates the dynamics of early and recycling endosomes as well as autophagosome-lysosomal fusion (De Matteis et al., 2017; Sharma et al., 2015). The protein has an N-terminal pleckstrin-homology (PH) domain (Mao et al., 2009), a central 5-phosphatase catalytic core (Tsujishita et al., 2001), a C-terminal ASH (ASPM–SPD2–Hydin) domain (Erdmann et al., 2007; McCrea et al., 2008), and a catalytically inactive RhoGAP (RhoGTPase activating protein)-like domain (Pirruccello and De Camilli, 2012). The C-terminal RhoGAP-like domain interacts with Rho family GTPases allowing recruitment to actin-rich membrane regions (Faucherre et al., 2005; Faucherre et al., 2003). The ASH/RhoGAP domain of OCRL interacts with the endocytic proteins APPL1 and Ses1 (also called IPIP27), associated with endocytosis and receptor recycling, respectively (Diggins and Webb, 2017; Noakes et al., 2011; Swan et al., 2010). Among the proteins that interact with OCRL, the Rab GTPases, which bind to the ASH domain, are most numerous. Interactions with Rab5 and Rab6 target OCRL to endosomes and the TGN (trans-Golgi network), respectively (Hyvola et al., 2006). Loss of OCRL function increases the amount of PI(4,5)P_2_ on endosomes impairing membrane trafficking events such as endocytosis/recycling of multiple classes of receptors and M6PR retrograde trafficking (Vicinanza et al., 2011).

Here we demonstrate that SdhA prevents endocytic and recycling vesicles from merging with the LCV by targeting OCRL. We found that *sdhA* mutants accumulate high levels of endocytic/recycling vesicles on the vacuole in an OCRL-dependent manner. In the process, SdhA interrupts OCRL interactions with Rab GTPases.

## Results

### SdhA contains multiple eukaryotic protein binding motifs

In search of host targets of SdhA, we found that its amino acid sequence predicted that the protein was connected to control of host cell endocytic dynamics. Sequence analysis found two putative “clathrin box” consensus sequences (Table S1), but also multiple endocytic sorting motifs predicted to bind adaptor complexes AP1, AP2, and AP3 (Table S1) (Edeling et al., 2006), as well as an OCRL-binding F&H motif (FxxxHxxØ) (Ø-bulky hydrophobic). This OCRL-binding motif is found in other endocytosis-associated proteins such as APPL1, Ses1, and Ses2 (latter two also called IPIP27A and IPIP27B; Fig. 1A) (Swan et al., 2010; Erdmann et al., 2007). Interestingly, SdhA and OCRL have similar arrangements of motifs predicting endocytic pathway association (Ungewickell et al., 2004; Mao et al., 2009). Given the presence of a potential OCRL binding site and the presence of an array of sites in SdhA that would direct it towards endocytic transport intermediates, we reasoned that SdhA might associate with OCRL (Fig. 1A). Such association could modulate endocytic processes that threaten the integrity of the *Legionella-*containing vacuole (LCV) (Anand et al., 2020).

**Figure 1.**
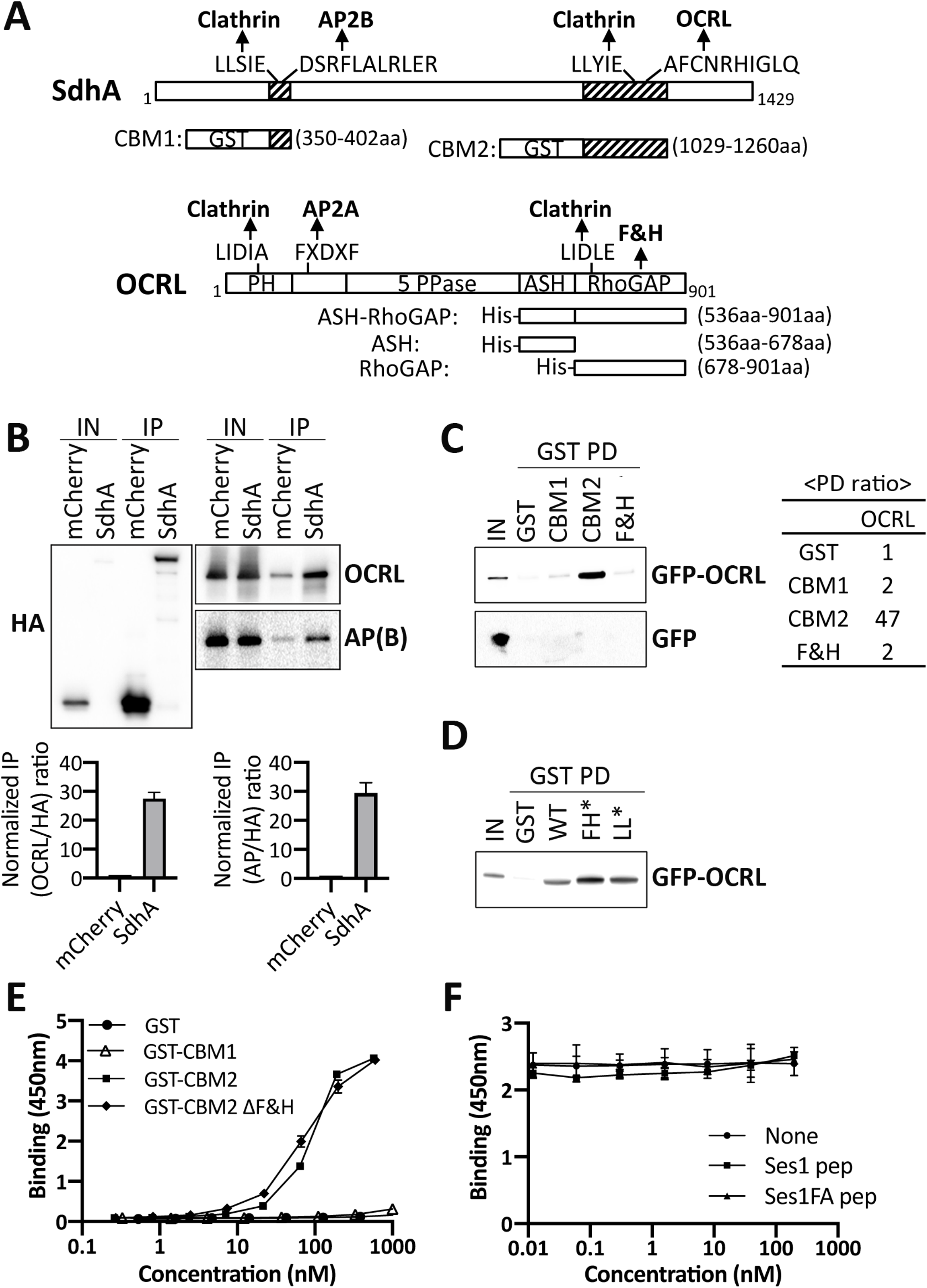
SdhA directly interacts with OCRL, but independently of F&H motif. (A) Conserved motifs of SdhA and OCRL, and maps of GST-or His-fusions. Motifs identified by Eukaryotic Linear Motif resource (www.ELM.eu.org). PH: pleckstrin homology; 5 PPase: 5 inositol polyphosphate phosphatase; ASH: ASPM-SPD2-Hydin; RhoGAP: Rho GTPase activating protein. (B) TOP: HA-mcherry or HA-mcherry-SdhA overexpressed in HEK cells were immunoprecipitated (IP) with anti-HA. AP(B): AP complex β subunit. The amount of input (0.2% lysates) and IP (20%) is shown. BOTTOM: Densitometry of co-immunoprecipitation. Average of two sets of independent experiments. (C) Purified GST fusions were used in pulldowns (GST PD) as described (Experimental Details). The amount of input (1% of total lysate) and resulting precipitate (20% of total) is shown. RIGHT PANEL: Ratio of the pull-down by densitometry. (D) GST-pulldowns as in (C) using SdhA-CBM2 mutations in F&H motif (F1195A H1199A; FH*) or clathrin box motif (L1177A L1178A; LL*). (E) 96-well plates coated with ASH/RhoGAP domain challenged with GST-tagged SdhA peptides (Experimental Details). SdhA binding to OCRL detected using anti-GST antibodies and chromogenic substrate (mean± SD, n=3). (F) Competition test of SdhA-CBM2 vs F&H motif peptide of Ses1 and OCRL binding-defective point mutation (Swan et al., 2010).

### SdhA interacts with OCRL

To determine if SdhA binds OCRL, we first performed co-immunoprecipitation (IP) with extracts of HEK cells transfected with HA-mCherry-tagged SdhA. The tagged SdhA construct quantitatively coprecipitated OCRL as well as the AP complex beta subunit when compared to mCherry alone, although the input level of HA-mCherry-SdhA was much lower relative to the mCherry-HA control (Fig. 1B). Normalizing for relative abundance of the HA tagged proteins present after immunoprecipitation compared to the input samples, association of OCRL with HA-mCherry-SdhA was approximately 27.5-fold above the control (Fig. 1B bottom left). SdhA also pulled down the AP complex beta subunit with similar efficiency (29.5-fold above control), albeit with higher nonspecific binding (Fig. 1B bottom right). Binding to clathrin was inconclusive due to nonspecific binding to mCherry (data not shown). Based on the strong interaction with OCRL, and potential for SdhA targeting an important regulatory protein involved in multiple endocytic paths, we dissected the interface between these two proteins further, and investigated its biological significance.

The interaction between SdhA and OCRL was tested using GST pull-down experiments. GST-tagged SdhA fragments were coupled to glutathione beads and incubated with extracts of HEK cells expressing either GFP-tagged full length OCRL or GFP. To this end, SdhA fragments containing either N-terminal motifs (Clathrin Box and AP complex binding Motif; CBM1) or C-terminal motifs (Clathrin Box and F&H Motif; CBM2) were tested for binding (Fig. 1A; CBM1, 350-402aa; CBM2, 1029-1260aa). The C-terminal GST-SdhA-CBM2 fragment bound to GFP-OCRL, but not GFP (Fig. 1C). The GFP-OCRL association with SdhA-CBM2 was about 47-fold above the control GST protein. In contrast, SdhA-CBM1 showed no binding to OCRL. The conserved amino acid peptide containing F&H motif (FxxxHxxØ) in Ses or APPL1 was shown to be sufficient for OCRL binding (Swan et al., 2010). However, a GST fusion containing the 13 amino acid F&H motif of SdhA did not bind to OCRL (Fig. 1C) and mutations in the putative F&H motif in SdhA-CBM2 did not disrupt OCRL binding (Fig. 1D), indicating that OCRL binding by SdhA is independent of the F&H motif.

To discount indirect binding of OCRL to SdhA by a complex series of interactions, we carried out solid-phase binding assays with purified GST-SdhA fragments and the ASH/RhoGAP domain of OCRL (536aa-901aa), the binding region for most of the OCRL partners (Fig. 1A). Direct binding was monitored by incubating increasing amounts of SdhA fragments with plate-immobilized ASH/RhoGAP domain of OCRL, probing with anti-GST. SdhA-CBM2 showed concentration-dependent binding to immobilized ASH/RhoGAP (Fig. 1E). In contrast, neither GST nor GST-CBM1 exhibited binding to ASH/RhoGAP. As predicted from the pulldown assay, binding of a CBM2 ΔF&H motif mutant was equivalent to CBM2, indicating that other sequences are responsible for binding to OCRL, with EC_50_ = 77.10 nM and 87.43 nM for mutant and WT respectively (Fig. 1E). SdhA binding to OCRL was further tested by competition with the known OCRL-binding F&H motif (13mer) of Ses1 (Swan et al., 2010). Addition of Ses1 F&H motif failed to decrease the binding efficiency of SdhA-CBM2, further arguing that the F&H motif of SdhA is not responsible for binding OCRL (Fig. 1F).

### Mapping the sites responsible for binding of SdhA and OCRL

Since SdhA does not require the F&H motif for OCRL binding, we searched for the OCRL binding site in SdhA-CBM2. The secondary structure of SdhA-CBM2 is predicted to contain 4 coiled-coils by ncoils (Lupas et al., 1991; Fig. 2A). Based on this prediction, each of the four coils was individually fused to GST and tested for ASH/RhoGAP binding. SdhA-CBM2A, encompassing residues 1029-1080 showed strong binding with the others showing low, but detectable binding to ASH/RhoGAP (Fig. 2B). Moreover SdhA-CBM2A (EC_50_∼27.2 nM) showed approximately 3-fold higher binding compared to the full SdhA-CBM2 fragment (EC_50_∼80.7 nM) consistent with other regions in SdhA-CBM2 interfering with SdhA-CBM2A binding to OCRL (Fig. 2C). The insolubility of SdhA precluded our ability to purify sufficient quantities to directly test if deletions in the full-length protein blocked binding to OCRL.

**Figure 2.**
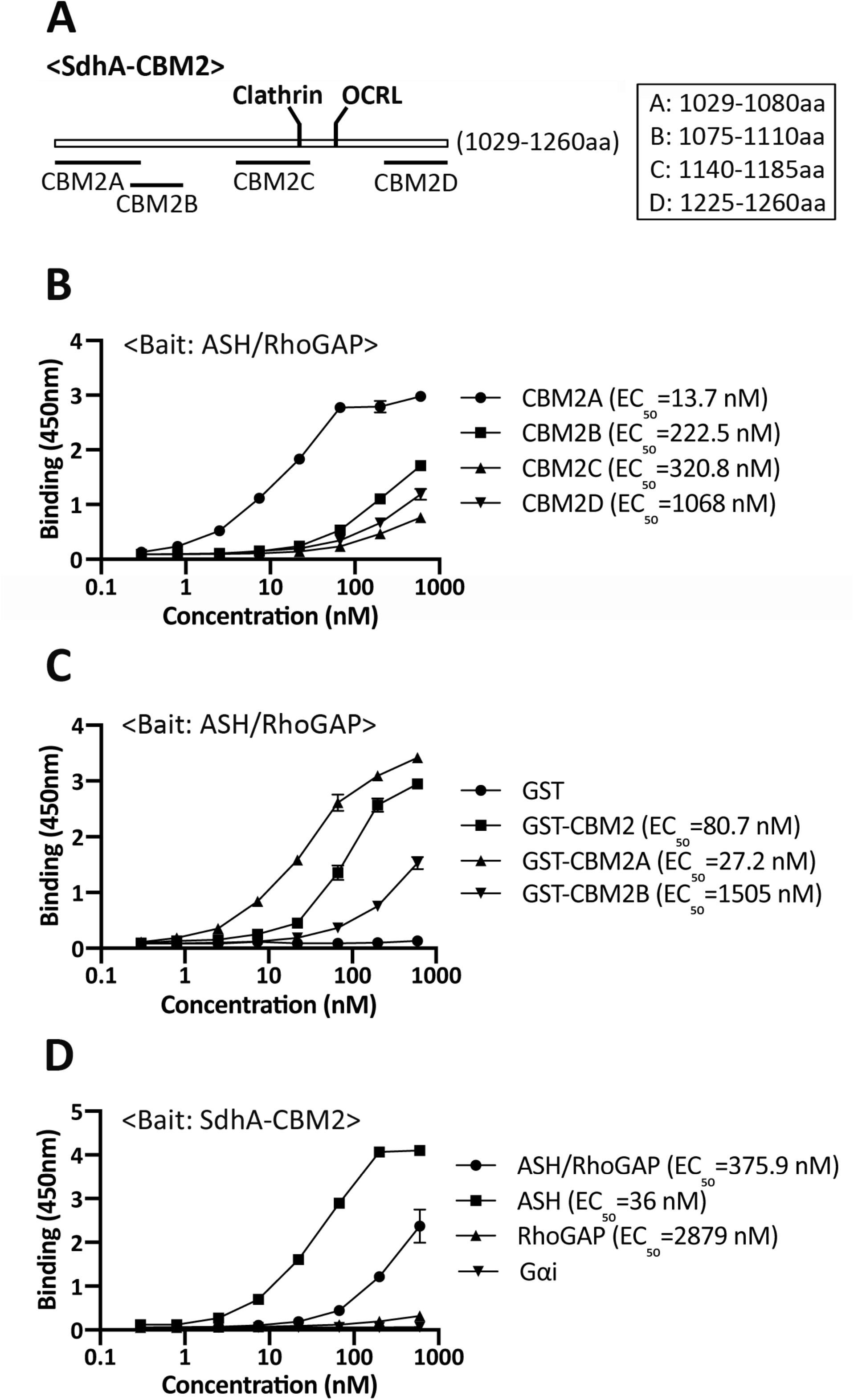
Mapping of the interacting regions of SdhA and OCRL using solid phase binding assays. (A)Constructs designed based on the coiled-coils prediction by ncoils (Lupas et al., 1991). (B) SdhA-CBM2A binds to ASH-RhoGAP domain. Plate coated with ASH/RhoGAP was challenged with indicated SdhA constructs. EC_50_ was calculated as described (Experimental Details). (C) High affinity binding of CBM2A to ASH-RhoGAP. Protocol as in 2(B). (D) SdhA-CBM2 associates with ASH domain specifically. Plate coated with SdhA-CBM2 was challenged with His-tagged OCRL domains. Binding detected by anti-His antibodies. His-Gαi was used as a negative control.

The RhoGAP domain of OCRL is the target of binding the F&H motifs in Ses and APPL1 (Pirruccello et al., 2011). To test for binding to the complete CBM2 fragment, we separated the ASH domain from the RhoGAP domain, and subjected purified proteins to the solid phase assay. SdhA showed stronger binding to the ASH domain than to the RhoGAP/ASH derivative and, strikingly, there was no detectable binding to the RhoGAP domain (Fig. 2D). Thus, the SdhA-CBM2 fragment binds nearby to the Ses/APPL1 binding region, but not on overlapping sites.

### The OCRL binding region of SdhA is essential for maintaining LCV integrity and promoting intracellular growth

To address the functional importance of SdhA binding to OCRL, we tested for its role in maintaining LCV integrity during bacterial infection by introducing SdhA deletion derivatives on plasmids into a *L. pneumophila* Δ*sdhA* strain. The truncation mutants lacking either CMB2 or CBM2A (Fig. 2A) had no distinguishable effect on growth in culture (Fig. S1A) or on LCV localization based on immunoprobing with SdhA antibody of macrophages after 3hrs incubations (Fig. 3A). Based on Western blot analysis, the expression levels of SdhA mutants were reduced relative to the levels of overproduced plasmid-borne SdhA-FL during in *vitro* growth (Fig. S1B), so their relative localization properties during infection were measured by scoring SdhA positive-LCVs at 4hrs after infection (Fig. S1C). Approximately 90% of LCVs scored positively for SdhA-FL, but none with empty vector indicating the probing is specific. The mutant protein ΔCBM2A showed indistinguishable levels of LCV localization compared to the wild type, although in the ΔCBM2 mutant about 40% of LCVs were positive. In contrast, a WT strain showing endogenous level of SdhA (not overproduced) did not show sufficient expression to detect SdhA using antibody probing. Therefore, even the most poorly expressed plasmid-borne SdhA mutant resulted in levels of LCV localization that were higher than the endogenously-expressed protein.

**Figure 3.**
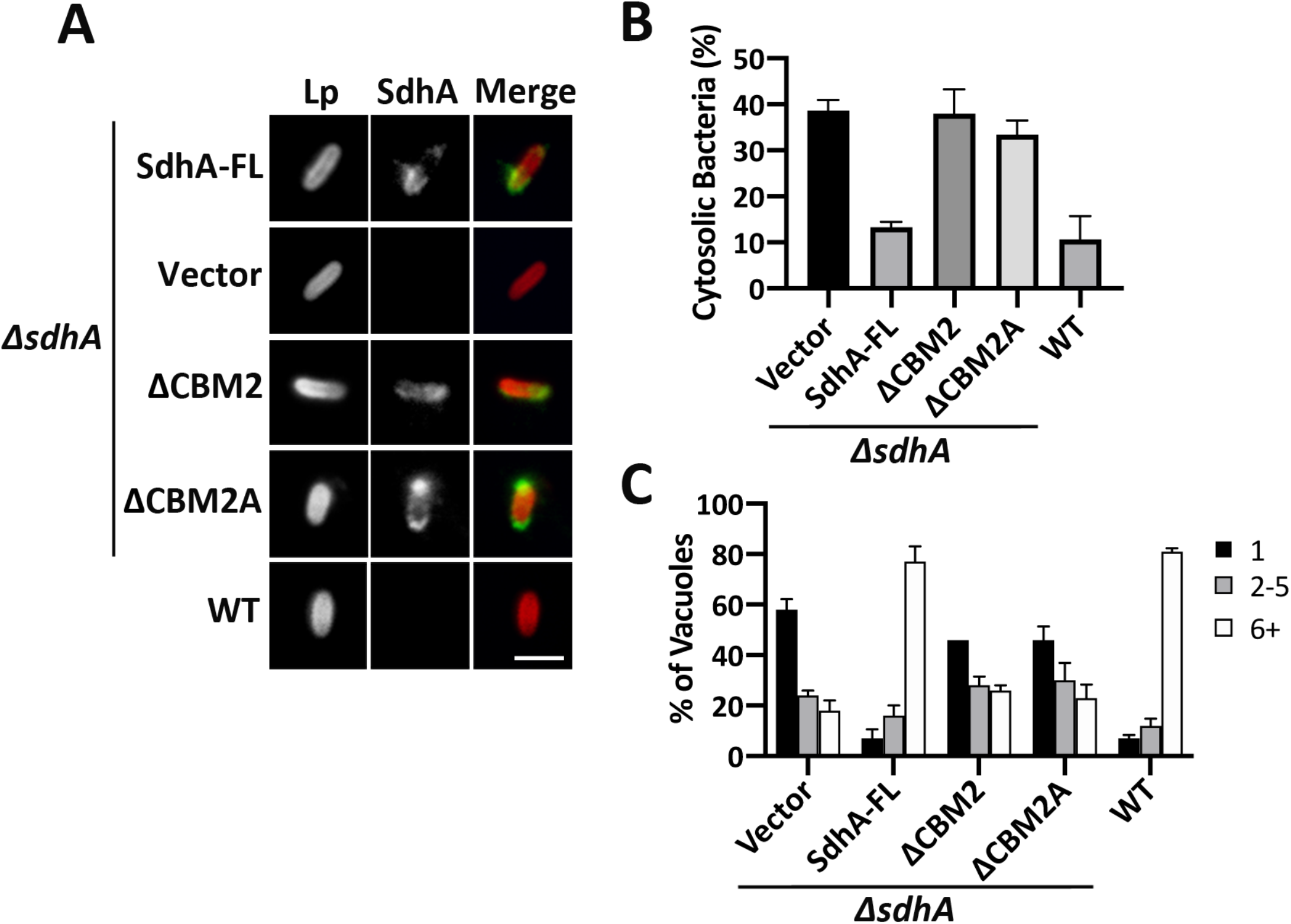
The OCRL binding region of SdhA is required for maintaining LCV integrity and intracellular growth in macrophages. BMDMs from the A/J mouse were challenged at MOI = 1 with *L. pneumophila* WT or *ΔsdhA* harboring SdhA variants. (A) SdhA localization on LCV was determined by immunofluorescence using antibodies against SdhA at 3hpi (Scale bar, 2 μm). See also Fig S1. (B) SdhA mutants do not rescue *ΔsdhA* vacuole disruption. Macrophages were challenged for 8hr, fixed, and stained for bacteria before and after permeabilization, and internalized bacteria in absence of permeabilization were quantified relative to total infected population (mean± SD, n=3). (C) Growth of *L. pneumophila* strains in macrophages, determined by the number of bacteria per vacuole 16h post uptake (mean± SD, n=3).

The mutants were next evaluated for vacuole disruption using our previously established immunofluorescence staining, based on antibody accessibility to bacteria in the absence of chemical permeabilization (Creasey and Isberg, 2012). By 8hrs after infection, about 40% of *ΔsdhA*-harboring vacuoles were permeable, compared to approximately 11% for the WT. Complementation of the *ΔsdhA* strain with FL-*sdhA* on the plasmid decreased the vacuole disruption to ∼10%, indistinguishable from WT strain Lp02 (Fig. 3B). In contrast, both ΔCBM2 and ΔCBM2A showed levels of LCV disruption that were similar to a Δ*sdhA* strain.

The loss of vacuole integrity triggered by a Δ*sdhA* strain causes a severe intracellular growth defect (Creasey and Isberg, 2012). As expected, we found that the internal deletions of *sdhA* resulted in growth defects that were indistinguishable from the total *sdhA* deletion strain (Fig. 3C). Taken together, these data are consistent with binding of OCRL by SdhA being tightly linked to maintaining LCV integrity.

### Ectopically expressed SdhA associates with OCRL-containing vesicles

To determine if SdhA-OCRL binding results in shared distribution through the cell, localization of the two proteins was examined by immunofluorescence microscopy in COS-7 cells transfected with mCherry-tagged SdhA and GFP-tagged OCRL. mCherry-SdhA was associated with membranous vesicles, mostly as giant ring-like structures (SdhA in Fig. 4A, SdhA-NOCO in Fig. S2), while mCherry alone localized to the nucleus and cytoplasm (Fig. 4A, CTR in Fig. S2). Surprisingly overexpression of SdhA dramatically altered the distribution of GFP-OCRL. SdhA overexpression caused OCRL aggregation around perinuclear region, with SdhA trapped within the OCRL-positive vacuoles, disturbing OCRL association with the Golgi and its normal association with punctate cytoplasmic vesicles (Fig. 4A, compare mCherry and SdhA). When we treated cells with the microtubule depolymerization drug nocodazole (Fig. S2, +NOCO), the aggregated OCRL-encompassed structures were distributed into small puncta that overlapped with SdhA signal, linking the aberrant morphology to microtubule function.

**Figure 4.**
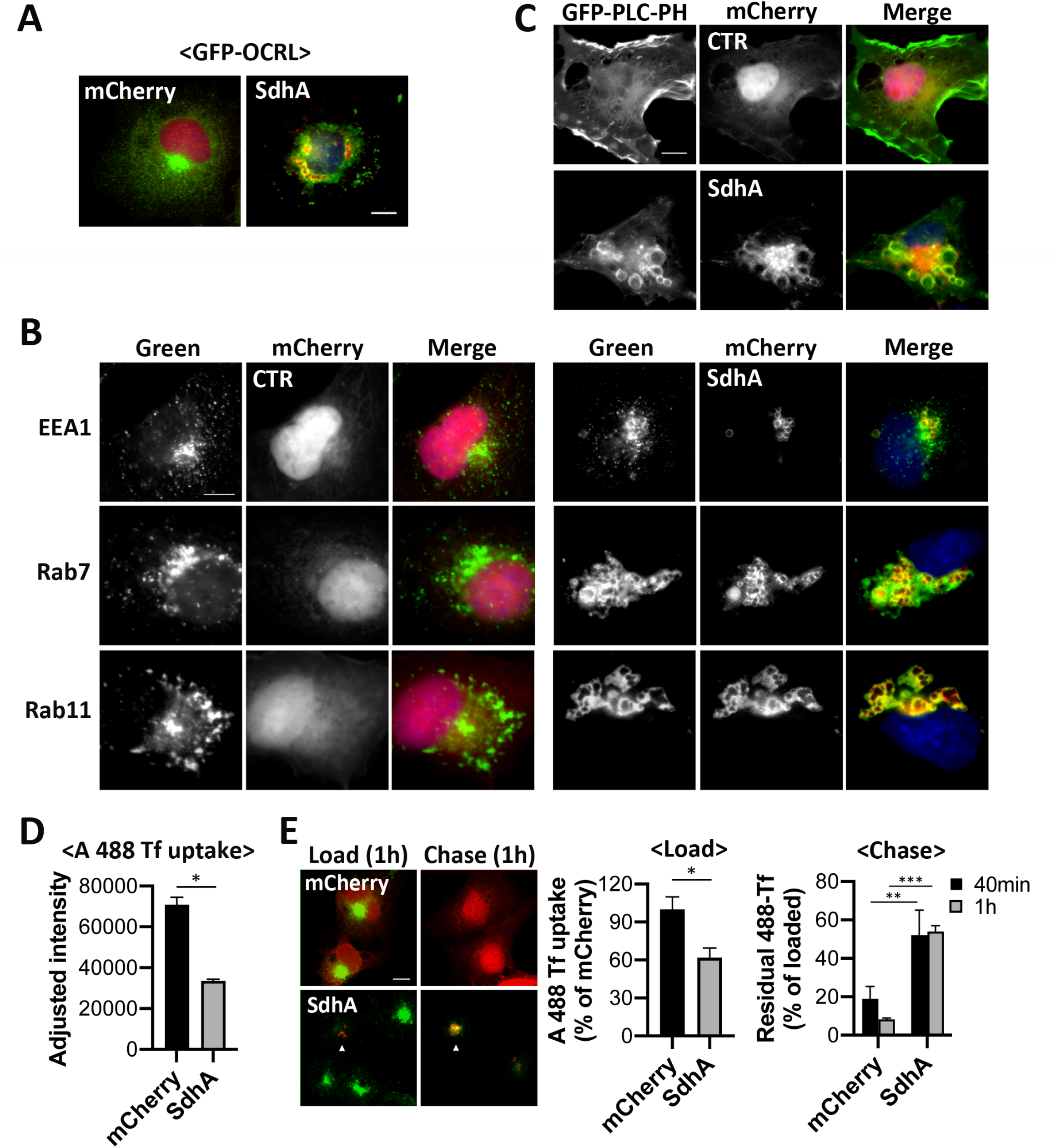
SdhA overexpression phenocopies loss of OCRL function. (A) Representative micrographs of fixed COS-7 cells co-expressing mCherry-SdhA (red) and GFP-OCRL (green). DNA labeled by Hoechst stain (blue). (Scale bar, 10 μm). See also Fig. S2. (B) mCherry-SdhA localizes to endocytic vesicles resulting in enlarged compartments. COS-7 cells expressing mCherry-SdhA (right panel) or mCherry (CTR, left panel) were either immunostained for EEA1 (green) or transfected with GFP-Rab7 or YFP-Rab11 (green). DNA labeled with Hoechst (blue). (C) COS-7 cells co-expressing mCherry-SdhA and GFP-PLCδ-PH. Additional images in Fig. S3. SdhA impairs endocytosis (D) and recycling of Tf (E). COS-7 cells were transfected with indicated expression vectors, mCherry or mCherry-SdhA. At 24 hr after transfection, cells were incubated with Alexa 488-Tf. (D) Uptake of Tf was measured as mean fluorescence intensities at 15 min. (E) For Tf recycling, the cells were loaded with Tf for 1 hr at 37°C (Load) and chased in complete medium for 40 and 60 min (Chase). Arrows indicate SdhA-transfected cells. The fluorescence intensity remaining in cell was quantified and expressed as percentages of the loaded Tf. Data are mean values ± SD (n=25 in triplicate) (*P<0.01; **P<0.05; ***P<0.001). Scale bars = 10 μm.

### SdhA overproduction phenocopies loss of OCRL function

To investigate whether the aberrant vacuoles generated by SdhA were the result of aggregated endosomes, we probed for endosomal markers in cells overexpressing mCherry-SdhA. Strikingly, SdhA-positive structures adopted unique morphologies, associating with endosomal markers of diverse origins, such as the early endosomal EEA1, late endosomal Rab7, and recycling endosome-derived Rab11 (Fig. 4B). In each case, SdhA redistributed into giant aggregated structures that were enveloped by a mixture of endosomal compartments. The abnormal vacuoles were strongly reminiscent of enlarged endosomal structures observed in cells defective for OCRL function (Vicinanza et al., 2011, Ben El Kadhi et al., 2011). Based on this result, we then determined if SdhA disrupts OCRL control of its preferential substrate, phosphatidylinositol-4,5-bisphosphate (PI(4,5)P_2_), and if it interferes with endosomal trafficking.

To this end, PI(4,5)P_2_ localization was probed using binding by GFP-PLCδ-PH. In control cells, PI(4,5)P_2_ associated with the plasma membrane (PM), particularly in regions of ruffling, as previously reported (Watt et al., 2002) (Fig. 4C). Remarkably, with SdhA-transfected cells, we found accumulation of PI(4,5)P_2_ on large vacuoles as well as depletion of PI(4,5)P2 in the PM, indicating dysfunctional PI(4,5)P_2_ homeostasis (Fig. 4C; Supplemental Fig. S3). The altered subcellular distribution of PI(4,5)P_2_ that we observed appeared to closely phenocopy previous observations in OCRL-depleted cultured mammalian cells, OCRL-depleted *Drosophila*, and OCRL deficient *zebrafish* (Vacinanza et al. 2011, Ben El Kadhi et al. 2011, Ramirez et al., 2012), consistent with SdhA overproduction interfering with OCRL function.

It has been reported that OCRL knockdown impairs uptake of transferrin (Tf) and slower recycling of internalized Tf from the PM (Vicinanza et al., 2011). Therefore, to probe for effects of SdhA on endosomal trafficking, recycling of transferrin receptor (TfR) was analyzed in cells overproducing SdhA by measuring internalization or recycling of Alexa488 (A488)-Tf. Compared with mCherry-transfected cells, transfection with SdhA showed a significant defect in uptake of Tf (Fig. 4D). The internalized pool of Tf also showed defective recycling to the PM, as Tf-preloaded cells chased for 1hr in the absence of probe lost approximately 50% of the accumulated Tf in SdhA-transfected cells, while 90% of the probe was lost from the control mCherry-transfected cells during the same time period (Fig. 4E). In addition, Tf accumulated in abnormal aggregated endosomes and was retained in clustered SdhA-containing structures, consistent with SdhA having direct disruptive effects on endosomal dynamics (Fig. 4E).

### SdhA inhibits Rab5 binding without interfering with the OCRL 5-phosphatase activity

We tested two models for how SdhA could antagonize OCRL function: altering OCRL association with target proteins or its catalytic function. The ASH domain of OCRL binds various Rab GTPases, most notably Rab5, Rab1, and Rab6 (Hyvola et al., 2006). As SdhA also binds to the ASH domain of OCRL, we tested if binding to SdhA fragments could block Rab5 association with OCRL. The binding affinities of constitutively active Rab5a (Q79L) and SdhA constructs were first tested, using the solid-phase binding assay in which the ASH/RhoGAP domain of OCRL was immobilized and incubated with increasing amounts of each protein (Fig. 5A). SdhA-CBM2 or SdhA-CBM2A exhibited a higher affinity for ASH/RhoGAP than Rab5a (Q79L). SdhA-CBM2A (EC_50_∼8.1 nM) showed approximately 12-fold higher binding compared to the Rab5a (Q79L) (EC_50_∼97.3 nM), arguing that SdhA may act by competing with known binding partners of OCRL (Fig. 5A). We further examined potential competition between SdhA fragments and Rab5a (Q79L) for OCRL binding. The ASH or ASH/RhoGAP domains were immobilized and challenged with 22 nM Rab5 in the presence of increasing amounts of SdhA-CBM2 or SdhA-CBM2A. Interestingly, the lower affinity SdhA-CBM2 fragment disrupted Rab5a (Q79L) binding to either the ASH or ASH/RhoGAP domains in a dose-dependent manner (Fig. 5B,C). Of note, the smaller fragment, SdhA-CBM2A, did not affect Rab5a (Q79L) binding to OCRL even though it has a higher apparent binding affinity for OCRL than SdhA-CBM2 (Fig. 2C). This is consistent with SdhA and Rab5 binding nonoverlapping sites on OCRL, with the larger CBM2 fragment blocking binding of Rab5 due to steric effects. As would be expected with the higher affinity interactions with SdhA, when either 22 nM of the CBM2 or 7.4 nM of CBM2A fragments were challenged for ASH or ASH/RhoGAP binding in the presence of increasing amounts of Rab5a (Q79L), the Rab protein had no effect on SdhA fragment binding to the OCRL domains (Fig. 5D,E). Thus, under the conditions tested here, the SdhA CBM2 fragment outcompetes Rab5 for binding to OCRL. This is consistent with our assays showing more efficient SdhA:OCRL binding than we observed for Rab5:OCRL interaction (Fig. 5A).

**Figure 5.**
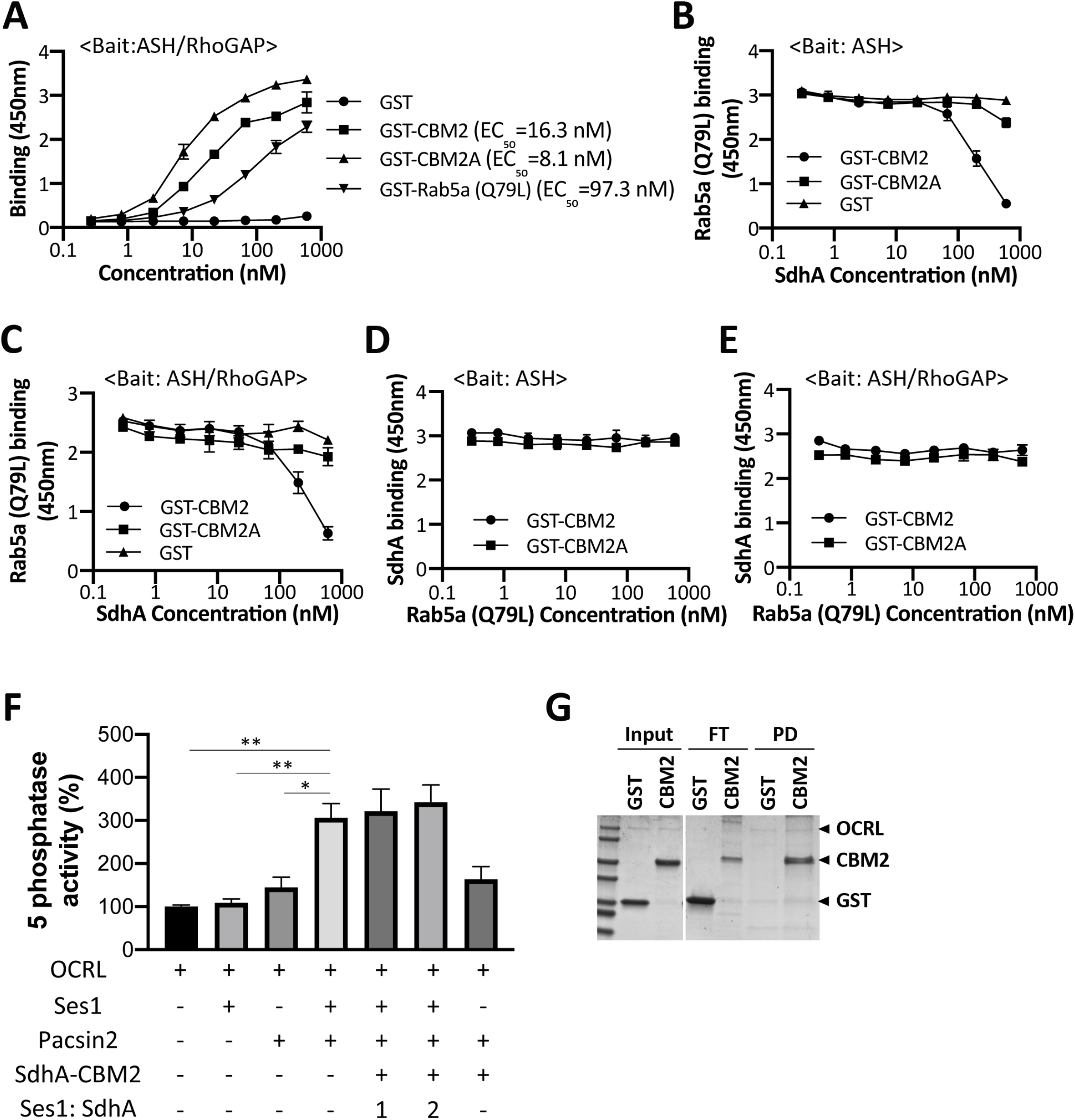
SdhA interrupts Rab5 binding. (A) High affinity binding of SdhA variants to OCRL. ASH-RhoGAP was immobilized and challenged with indicated GST-tagged proteins. EC_50_ were calculated and expressed as nM (Experimental Details). (B) Competitive binding of Rab5 and SdhA to ASH domain. Immobilized ASH domain was challenged with 22nM Rab5a (Q79L) in combination with increasing amounts of GST-fused SdhA constructs. (C) Same as in (B), except ASH/RhoGAP domain was immobilized. (D) SdhA binding to ASH domain of OCRL was not affected by challenge with Rab5a. Same as in (B), but constant amounts of SdhA (CBM2=22nM, CBM2A=7.4nM) and increasing amounts of Rab5a (Q79L). (E) Same as in (D), but ASH/RhoGAP domain was immobilized. (F) SdhA-CBM2 does not affect the Ses1- and Pacsin2-stimulated 5-phosphatase activity of OCRL. Phosphatase activity was measured as described (Experimental Details). His-OCRL (50 nM) was incubated with indicated proteins and with 200 μM of PI(4,5)P_2_-containing liposomes. Data expressed as percentage of 5-phosphatase activity compared to incubation of OCRL with liposomes. Data represent triplicate assays ±SE (standard error). (*P<0.01; **P<0.05; ***P<0.001) (G) SdhA-CBM2 binding to OCRL during 5-phosphatase assay in (F) demonstrated by pulldown assay. His-OCRL and SdhA-CBM2 were collected with Ni^2+^ resin and analyzed by Coomassie staining. The amount input (10%), unbound FT (flowthrough, 20%) and bound PD (pulldown 50%) are displayed.

We next analyzed whether SdhA directly affects the catalytic activity of OCRL as a consequence of binding the ASH domain, which is proximal to the 5-phosphatase domain of OCRL (Fig.1A). To this end, the 5-phosphatase activity was assayed using purified OCRL incubated with PI(4,5)P_2_-containing liposomes. The 5-phosphatase activity of OCRL is inherently weak in published assays in the absence of a source of stimulation, making inhibitory effects difficult to detect (Billcliff et al., 2015). It has been shown that the activity is stimulated by formation of tripartite complex with Ses1 and Pacsin2 (Billcliff et al., 2015), so we used these components to test if SdhA interferes with the 5-phosphatase activity. Consistent with previous results, the 5-phosphatase activity was dramatically stimulated in the presence of both Ses1 and Pacsin2. Addition of SdhA-CBM2 in the presence of this complete reaction mix, however, showed no significant depression of the stimulated activity (Fig. 5F). SdhA-CBM2 was clearly competent to bind OCRL using these assay conditions, as His-OCRL efficiently pulled down GST-SdhA-CBM2, but not GST alone (Fig. 5G). These results indicate that SdhA likely hijacks OCRL, interrupting binding of cellular partners without disrupting its phosphatase activity.

### Cellular depletion of OCRL enhances the integrity of LCVs harboring the *sdhA* mutant

To assess the role of OCRL in modulating LCV integrity, OCRL was efficiently RNAi-depleted in COS-7 cells (Fig. 6A). The OCRL-depleted cells were next challenged for either 4 or 8 hrs with Δ*sdhA* or WT strains and the relative levels of disrupted LCVs was determined by immunostaining (Experimental Procedures). Cells depleted of OCRL showed a significant enhancement in the integrity of vacuoles harboring the Δ*sdhA* strain at both time points, indicating that the absence of OCRL stabilized the *sdhA*-containing vacuole (Fig. 6B). OCRL knockdown also resulted in enhanced intracellular growth of Δ*sdhA* as well as WT in COS-7 cells (Fig. 6C).

**Figure 6.**
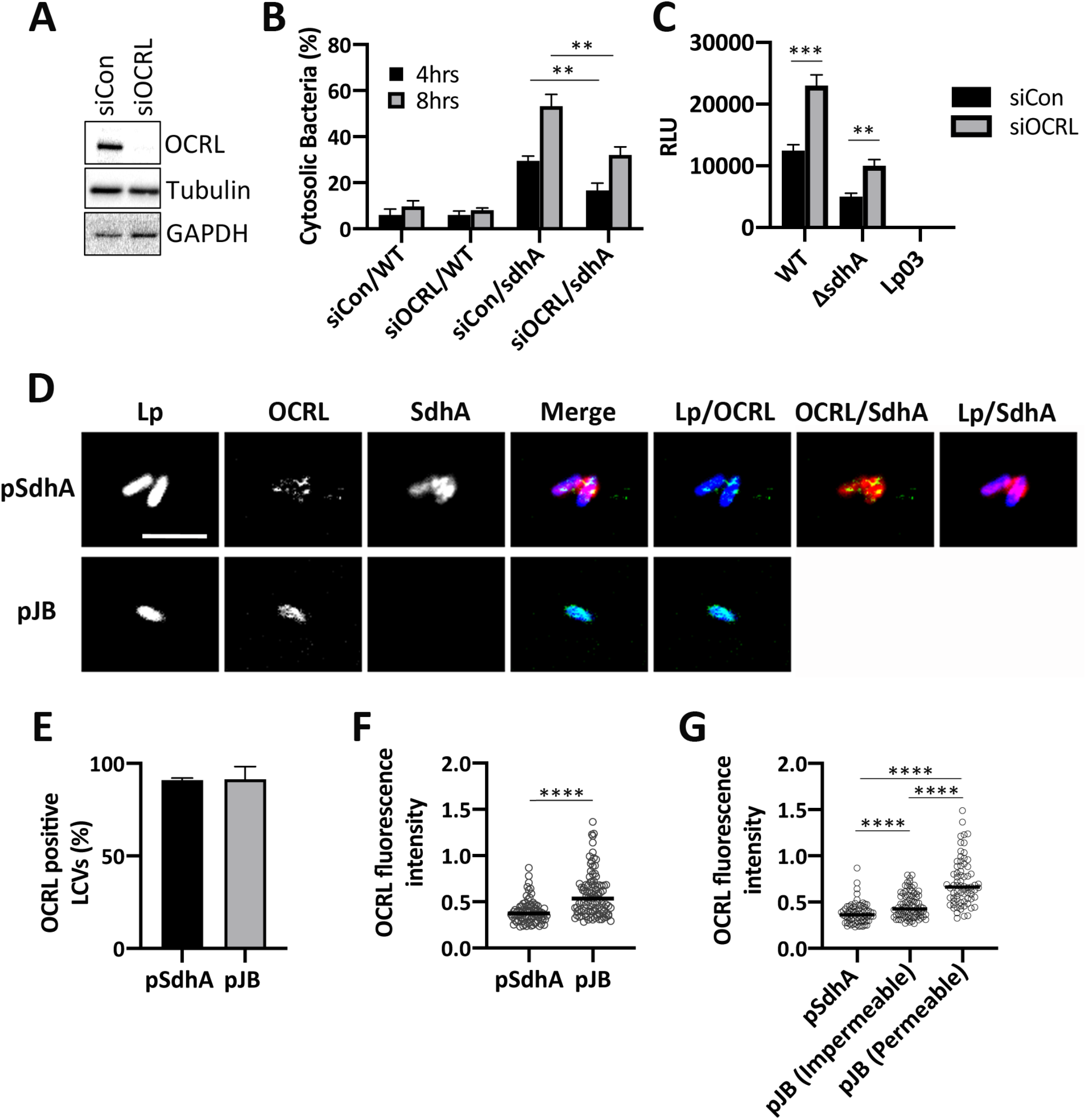
OCRL is linked to vacuole disruption of *L. pneumophila sdhA* mutants. (A) COS-7 cells treated with OCRL-targeting (siOCRL) or non-targeting control siRNA (siCon) for 72hrs were gel fractionated and immunoprobed with antibodies directed against noted proteins. (B) COS-7 cells were depleted by siRNA (3 days) and then challenged at MOI =5 with noted *L. pneumophila* strains. Cells were fixed and stained for bacteria before and after permeabilization, as in Fig. 4B. Data are mean values ± SD (n=3) (**P<0.05). (C) *L. pneumophila lux*^*+*^ strains were incubated with COS-7 cells at MOI = 20 and bacterial yield was measured 48 hrs post infection by relative luminescence (RLU). The replication-deficient *dotA* null mutant Lp03 was used as a negative control. Data are mean values ± SD (n=3). (D) Confocal image showing a section of vacuoles isolated from infected U937 cells (MOI = 10, 3h) with Δ*sdhA* mutant harboring pSdhA or pJB vector. The presence of OCRL and SdhA on LCVs was assessed by immunofluorescence using antibodies directed against OCRL, SdhA and *L. pneumophila*. (Scale bar, 4 μm). See also Fig S4. (E) Quantification of OCRL positive LCVs containing Δ*sdhA* mutants with pSdhA or pJB vector at 3hpi. Data are mean values ± SD (n=3) (>85 LCVs each replicate). (F) Plot of OCRL intensity associated with LCV (n>70), with medians displayed. (G) Plot comparing the OCRL intensity on LCVs harboring pSdhA compared to strain harboring empty pJB vector, divided into impermeable and permeable LCVs (n>70). (****P<0.0001, Mann-Whitney U-test)

### OCRL accumulates on the disrupted vacuole of *sdhA* mutants

Previous work has shown that OCRL localizes to LCVs in *D. discoideum* and RAW264.7 macrophages (Weber et al., 2009). To determine if SdhA is involved in the localization of OCRL on LCVs, we challenged U937 macrophages for 3hrs with a Δ*sdhA* strain harboring either a plasmid overproducing SdhA or an empty vector control. After homogenization of infected macrophages, the localization of SdhA and endogenous OCRL on LCVs in post-nuclear supernatants was determined by immunofluorescence microscopy. Approximately 90% of the LCVs stained positively for OCRL regardless of the presence of SdhA (Fig. 6D,E). The pattern of OCRL localization on the LCVs, however, could clearly be differentiated between the two strains. In the presence of SdhA, OCRL formed punctate structures associated with LCVs that appeared to be enveloped by SdhA. In contrast, there was dense circumferential accumulation of OCRL about the LCV in absence of *sdhA* (Fig. 6D) (more images in Fig. S4). Based on image analysis, the median fluorescence intensity of OCRL accumulation about vacuoles was approximately 1.4 times greater in the absence of SdhA than in its presence, with a broad distribution of intensities observed for the mutant, pointing toward two populations of LCVs (P<0.0001; Mann-Whitney U-test; Fig. 6F). To investigate this distribution further and test if vacuole disruption was related to accumulated OCRL, we quantified OCRL on intact or disrupted LCVs (Fig. 6G). There was a significant correlation between OCRL intensity and vacuole disruption. For the strain lacking SdhA, the median OCRL fluorescence intensity of disrupted LCVs was 1.6-fold greater compared to those with intact LCVs (P<0.0001; Mann-Whitney U-test). The OCRL intensity of intact LCVs containing the empty vector strain was significantly higher than the OCRL intensity of LCVs containing the pSdhA strains, indicating OCRL accumulation occurs before vacuole disruption and then continues to accumulate (Fig. 6G). Therefore, SdhA interferes with the accumulation of OCRL on LCVs and is associated with vacuole disruption.

### SdhA inhibits the accumulation of OCRL-dependent endosomal traffic to LCVs

The localization of several well-characterized host proteins associated with OCRL-controlled endosomal traffic was assessed by immunofluorescence microscopy of LCVs from postnuclear supernatants of U937 cells (Vicinanza et al., 2011). We quantified the number of LCVs positive for endosomal EEA1, retrograde cargo trafficking cation-independent mannose 6-phosphate receptor (CIMPR), and TfR, linked to recycling cargo (Fig. 7A). For EEA1 and CIMPR, there was a significant increase in LCV association for each marker after infection with a Δ*sdhA* strain for 3hrs when compared to WT. The increase was particularly striking with EEA1, supporting our previous analysis of the behavior of Δ*sdhA* strains with murine bone marrow-derived macrophages (Anand et al., 2020) (Fig. 7A).

**Figure 7.**
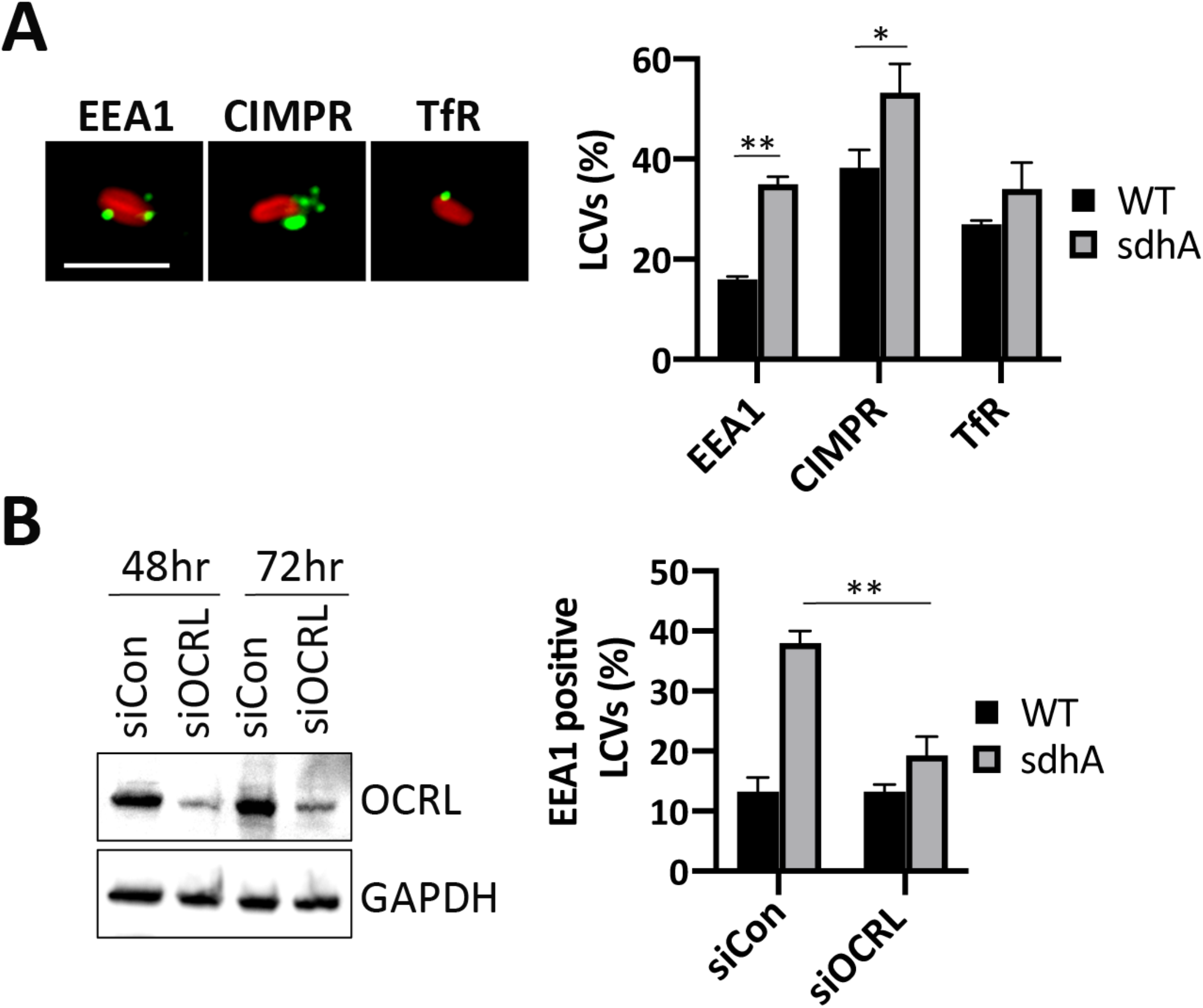
OCRL-dependent accumulation of endosomal compartments on vacuole surrounding *ΔsdhA* mutant. (A) U937 macrophages were challenged for 3 hr with L. *pneumophila* wild type and *ΔsdhA* (MOI = 10). The presence of endosomal EEA1, retrograde trafficking cargo CIMPR, and recycling endosomal TfR on LCVs was evaluated by immunofluorescence microscopy in postnuclear supernatants of infected cells (Experimental Details). Data are mean values ± SD of triplicate samples. Scale bar represents 4 μm. (B) LEFT PANEL: Effect of siOCRL on protein expression in U937 cells. GAPDH was used as a loading control. Gel fractionated samples were immunoprobed with indicated antibodies. RIGHT PANEL: The presence of EEA1 on LCVs was quantified from the infected U937 cell lysates as in panel (A). (*P<0.01; **P<0.05)

To assess whether enhanced accumulation of EEA1 on LCVs harboring *sdhA* mutants is dependent on OCRL, OCRL was depleted by siRNA prior to infection. In comparison to the control with scrambled siRNA, EEA1 positive LCVs harboring the Δ*sdhA* strain decreased to a level that was indistinguishable from WT (Fig. 7B). This indicates that aberrant trafficking of EEA1 to LCVs harboring the Δ*sdhA* mutant appeared largely dependent on OCRL function, consistent with SdhA controlling OCRL function for the purpose of blockading early endosomal vesicles.

## Discussion

Mammalian OCRL protein and its *Dictyostelium discoideum* homologue Dd5P4 localize to the *Legionella-*containing vacuole (LCV), limiting intracellular replication of *L. pneumophila* (Weber et al., 2009, Finselet at al., 2013). In our study, we argue that OCRL promotes events that disrupt LCV integrity, with SdhA protein being a key player in preventing OCRL restriction of pathogen growth. As SdhA is one of the few Icm/Dot translocated substrates required for survival in primary macrophages, the interface of SdhA with this host inositol polyphosphate 5-phosphatase is likely to be a critical step in controlling the balance between host innate restriction and proliferation of the pathogen. The OCRL phosphatase activity is known to control a number of membrane trafficking steps. The results documented here argue that SdhA blocks an OCRL-regulated step in the movement of vesicles from an endosomal compartment, resulting in disruption of the LCV membrane. Presumably the blockade occurs by SdhA hijacking of OCRL.

We have found that RNAi depletion of OCRL in cells challenged with the *sdhA* mutant significantly increases the number of intact LCVs, consistent with a role for OCRL in driving vacuole membrane disruption. Similarly, OCRL depletion in COS-7 cells stimulates intracellular replication of the *L. pneumophila* Δ*sdhA* (Fig. 6B,C). Immunofluorescence detection of OCRL interaction with the LCV indicates that there are likely two modes of OCRL interface with the vacuole (Fig. 6). On encounter with vacuoles harboring WT *L. pneumophila*, OCRL-staining compartments target to SdhA-rich regions. In contrast, loss of SdhA function enhances OCRL circumferential recruitment about the LCV, and this recruitment appears particularly robust in vacuoles undergoing disruption (Fig. 6G). Taken together, these results argue that the LCV surrounding the *sdhA* mutant vacuole merges with OCRL-containing compartments, resulting in eventual vacuole disruption. In cells harboring the WT strain, vacuole integrity is maintained as a consequence of diversion of these compartments by SdhA.

In previous work, we presented evidence that SdhA likely acts by preventing access of endosome-derived compartments to the LCV (Anand et al., 2020). This model was based on RNAi screens demonstrating that disruption of Δ*sdhA* mutant-containing vacuoles can be reversed by depletion of endocytic Rab GTPases. A primary consequence of SdhA loss was shown to be the accumulation of early endosomal protein EEA1 on the defective vacuole in primary macrophages, dependent on the function of Rab5 and Rab11 (Anand et al., 2020). As a number of GTPases involved in endosome dynamics, such as Rab5 and Rab8 are known interacting partners of OCRL (Grant and Donaldson, 2009), we asked whether disrupting OCRL function similarly could block this aberrant EEA1 accumulation. The depletion of OCRL dramatically decreased EEA1 association with the Δ*sdhA* vacuole (Fig. 7B).

Based on these results, we hypothesize that in the absence of SdhA, OCRL and Rab GTPase interactions recruit endosomal compartments to the Δ*sdhA* vacuole and disrupt the pathogen niche. In the vacuole harboring WT bacteria, SdhA can bind OCRL and interfere with Rab binding to the phosphatase, preventing OCRL-containing compartments from directly targeting the vacuolar membrane. The fact that SdhA binds to the ASH domain of OCRL (Fig. 2) and sterically blocks Rab protein interaction with OCRL may be significant in this regard. OCRL mutants defective in Rab binding have been shown to result in aberrant OCRL targeting, and depletions of Rab1 or Rab6 show similar defects (Hyvola et al. 2005). Therefore, association of OCRL-containing vesicles with SdhA and consequent disruption of Rab protein binding could prevent Rab effectors from promoting efficient docking of either OCRL or its associated endosomes with the LCV surface.

The relatively small CBM2A region in the C-terminal of SdhA (aa1029-80), predicted to form one of several coiled-coil structures throughout the protein, is sufficient to bind the OCRL ASH domain (Fig. 2). This region in SdhA is functionally important, as overproduction of a variant of the protein that precisely deletes this region fails to complement the Δ*sdhA* mutation (Fig. 3 and Fig. S1). Consistent with the defect being tightly associated with loss of OCRL control, OCRL is profoundly altered in transfectants overexpressing SdhA, resulting in giant OCRL-encompassed vacuoles that surround SdhA-rich regions (Fig. 4). Loss of OCRL function has been documented to generate large vacuoles harboring endosomal components, thought to result from blockading of endosomal traffic to the Golgi (Vicinanza et al., 2011, Choudhury et a., 2005, Kadhi et al., 2011). As a consequence, disrupted OCRL causes accumulation of PI(4,5)P_2_ in endosomal compartments and disruption of transferrin receptor (TfR) recycling (Vicinanza et al., 2011, Choudhury et a., 2005, Kadhi et al., 2011). Overexpression of SdhA exactly phenocopies the functional loss of OCRL, as we have demonstrated that SdhA transfectants cause both mislocalization of PI(4,5)P_2_ and dysfunctional TfR recycling (Fig. 4).

It seems counterintuitive that OCRL function should be associated with LCV disruption. The LCV is rich in PI4P, so it might be thought that OCRL would play a collaborative role in LCV biogenesis, as its inositol polyphosphate 5-phosphatase activity can generate PI4P, which in turn anchors a wide swath of Icm/Dot effector proteins to the LCV cytoplasmic surface (Weber et al., 2006; (Hsu et al., 2012)). *L. pneumophila* has a number of well-characterized translocated inositol phosphate kinase and phosphatases capable of modulating PI4P dynamics, consistent with maintaining PI4P homeostasis in the LCV (Dong et al., 2016; Hsu et al., 2012; Toulabi et al., 2013). It seems likely that OCRL plays a surprising negative role by stimulating the biogenesis, recruitment or docking of forbidden membrane compartments that act to destabilize the LCV rather than directly modifying the LCV. Alternatively, the presence of OCRL on the vacuolar membrane could disrupt the PI4P homeostatic balance provided by *Legionella* Icm/Dot effectors, resulting in overload of this lipid and hypersensitivity to phospholipases.

As we have previously argued, SdhA belongs to a larger class of pathogen proteins called vacuole guards that act to prevent intravacuolar microbial pathogens from being attacked by disruptive membrane components largely derived from endosomes and recycling compartments (Anand et al., 2020). SdhA fits in well with this class of proteins, as blocking vesicular transit of these compartments to the LCV increases vacuole stability (Anand et al., 2020). By binding OCRL, SdhA may play a role in modulating self-nonself recognition by the LCV. Forbidden compartments harboring OCRL may have membrane compositions rich in PI4P that are similar to that of the LCV. This, in turn, could direct targeting and fusion of these disruptive compartments with the LCV. SdhA acts to “guard” the pathogen-containing vacuole by binding and diverting OCRL, preventing either direct interaction with disruptive compartments or blocking association of OCRL with the LCV.

Although the newly described SdhA biochemical activity has not been observed previously, the connection of OCRL to the endocytic and recycling compartments increases documented parallels between SdhA and the *Salmonella* SifA protein (Beuzon et al., 2000; McGourty et al., 2012). Both SifA and SdhA are required to maintain the integrity of their respective vacuoles, with failure to prevent host cell-mediated disruption resulting in release of bacteria into the cytosol and activation of a Caspase 4/11-Gasdermin D-dependent pyroptotic response in phagocytes (Aachoui et al., 2013; Casson et al., 2015; Pilla et al., 2014; Shi et al., 2015). An intimate connection between maintaining vacuole integrity and preserving appropriate lipid content of the vacuolar membrane has long been suspected in both cases, primarily based on suppressor mutation analysis (Creasey and Isberg, 2012; Kolodziejek et al., 2019). Finally, our work argues that SdhA regulates the host cell endocytic/recycling pathways, reminiscent of the demonstrated role of SifA in hijacking retrograde cellular transit and controlling egress of CIMPR (Dumont et al., 2010; McEwan et al., 2015; McGourty et al., 2012), as observed here (Figs. 4E, 7A). This argues that components of the endosomal pathway act as important disruptive forces that can block pathogen growth without directly targeting the organism into a lysosomal compartment. The details of the nature of these disruptive forces, and the phospholipid composition that results in destabilization of these compartments, remain to be determined.

## Acknowledgements

This work was supported by HHMI and NIAID grants R01 AI113211 and R01 AI146245 to RRI. We thank Ila Anand for scientific discussions throughout the course of this work, and Kristen Davis, Wenwen Huo, Erion Lipo, and Seongok Kim for review of the manuscript. We thank Dr. Pietro De Camilli for providing OCRL antibody and GFP-OCRL construct; Dr. Matthias Machner for Rab5a (Q79L) construct. We also thank Drs. Elizabeth Draganova and Ellen White for patient help setting up the lipid extrusion assays and with Baculovirus expression.

---

**Figure S1.**
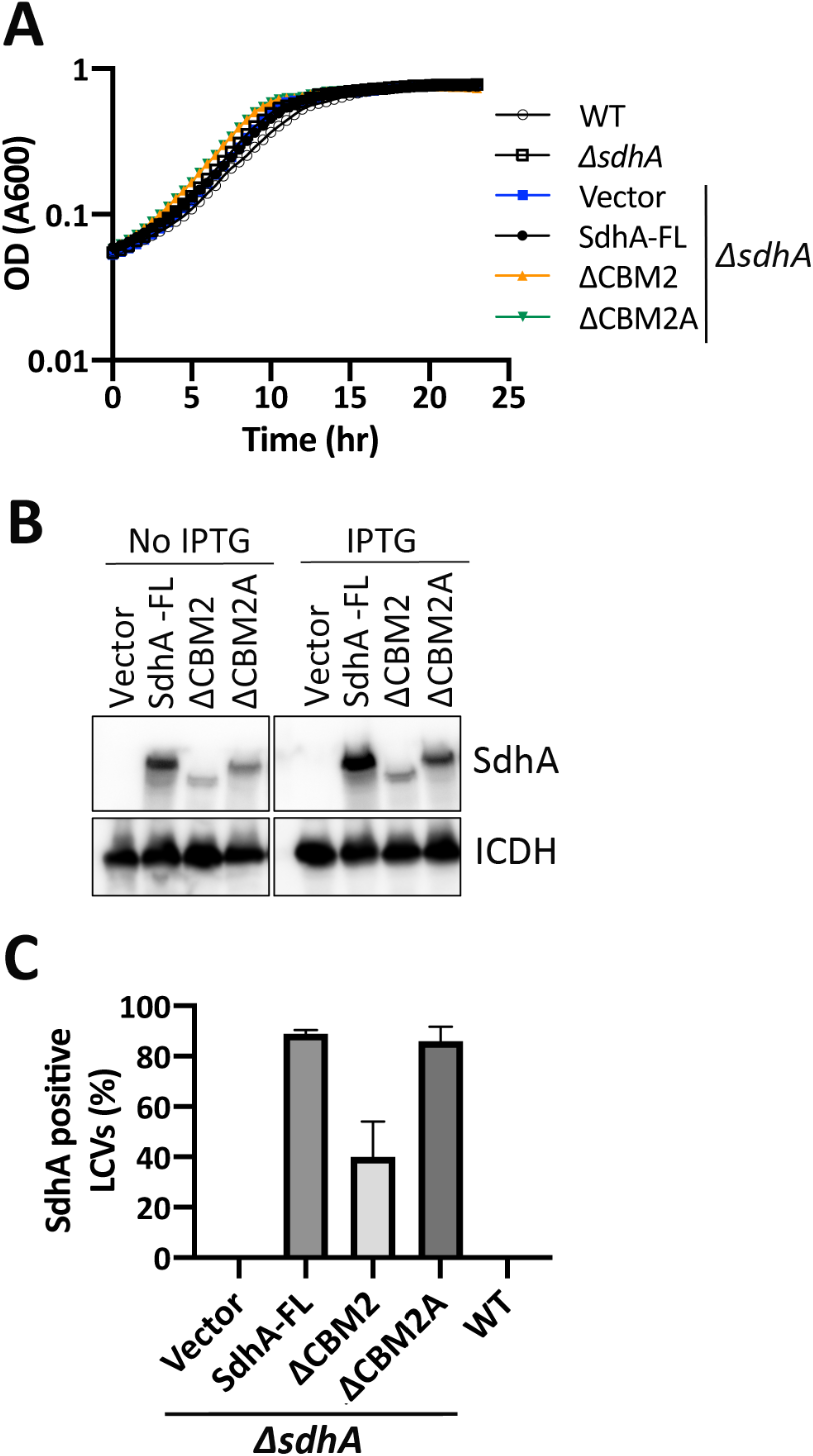
Characterization of SdhA mutants. (A) Growth of noted strains in AYE broth. Strains with *ΔsdhA* allele have noted genes inserted into pJB908 (Vector). (B) Western blot analysis of expression levels of SdhA variants. Isocitrate dehydrogenase (ICDH) was used for loading control. (C) The presence of SdhA variants on LCV surface was assessed by fluorescence microscopy as in Fig. 3A. SdhA is undetectable in WT strain and must be overproduced on pJB908 to identify by immunofluorescence microscopy. Each of the *ΔsdhA* strains harbor pJB908 with noted chromosomal fragments. Data represents means and SDs of triplicates.

**Figure S2.**
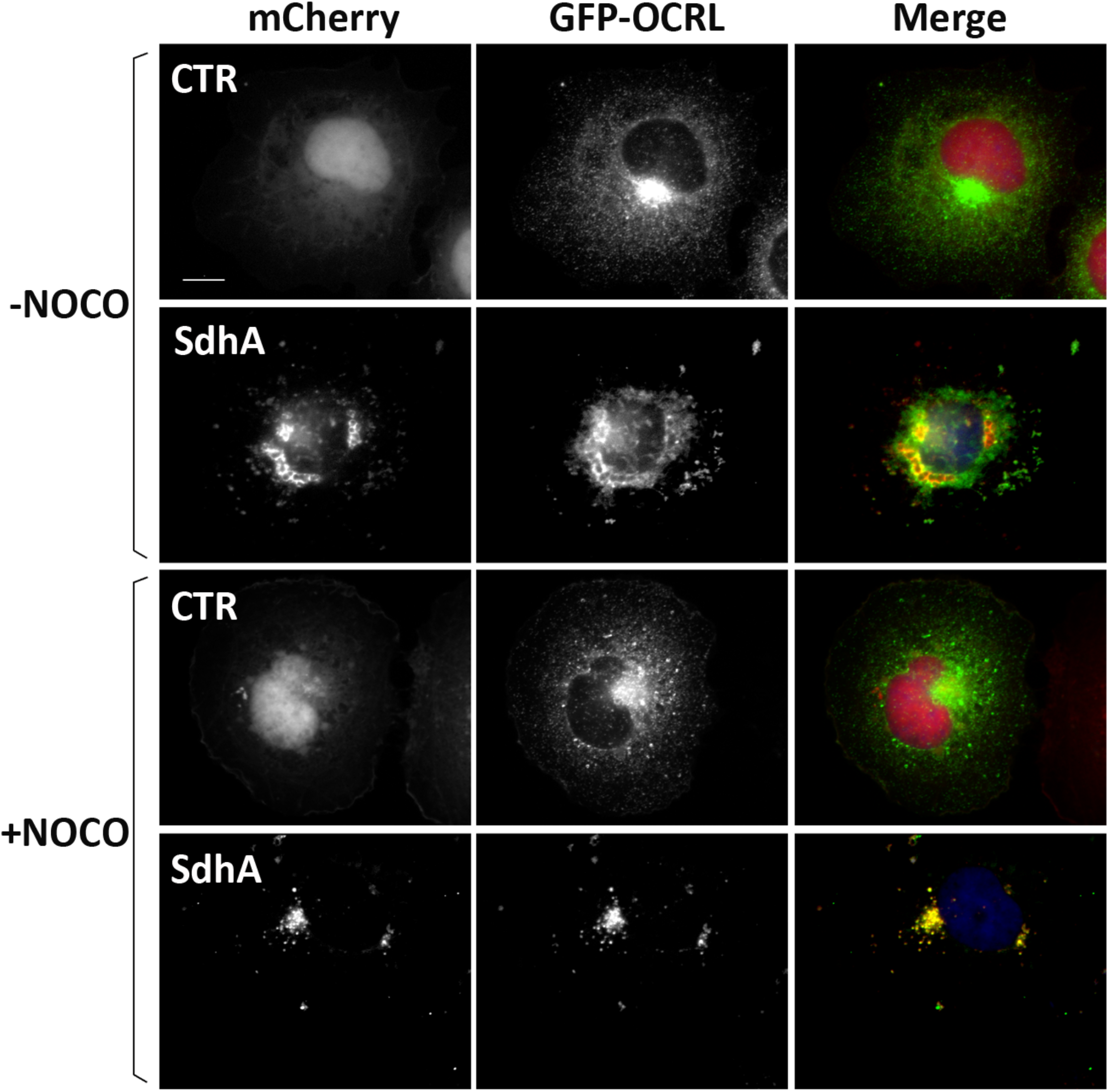
Ectopically expressed SdhA associates with OCRL positive compartments. Representative micrographs of fixed COS-7 cells coexpressing mCherry-SdhA (red) and GFP-tagged OCRL. DNA was labeled by Hoechst stains (blue). Cells were treated with nocodazole (NOCO) to release aggregation of the compartments. (Scale bar, 10 μm)

**Figure S3.**
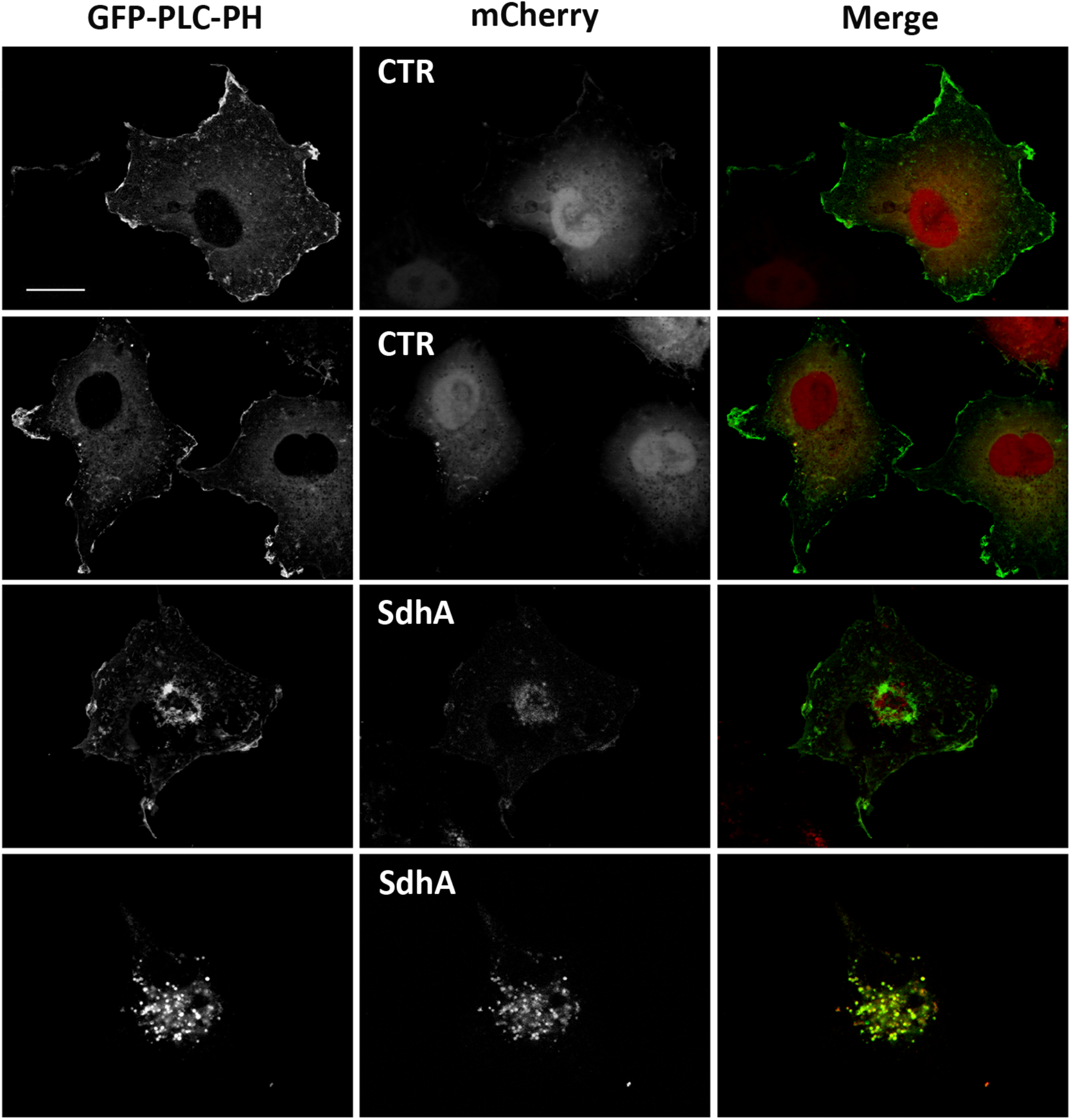
COS-7 cells coexpressing mCherry-SdhA and GFP-PLCδ-PH demonstrate rearrangement and internalization of PI(4,5)P_2_. Scale bar represents 20 μm.

**Figure S4.**
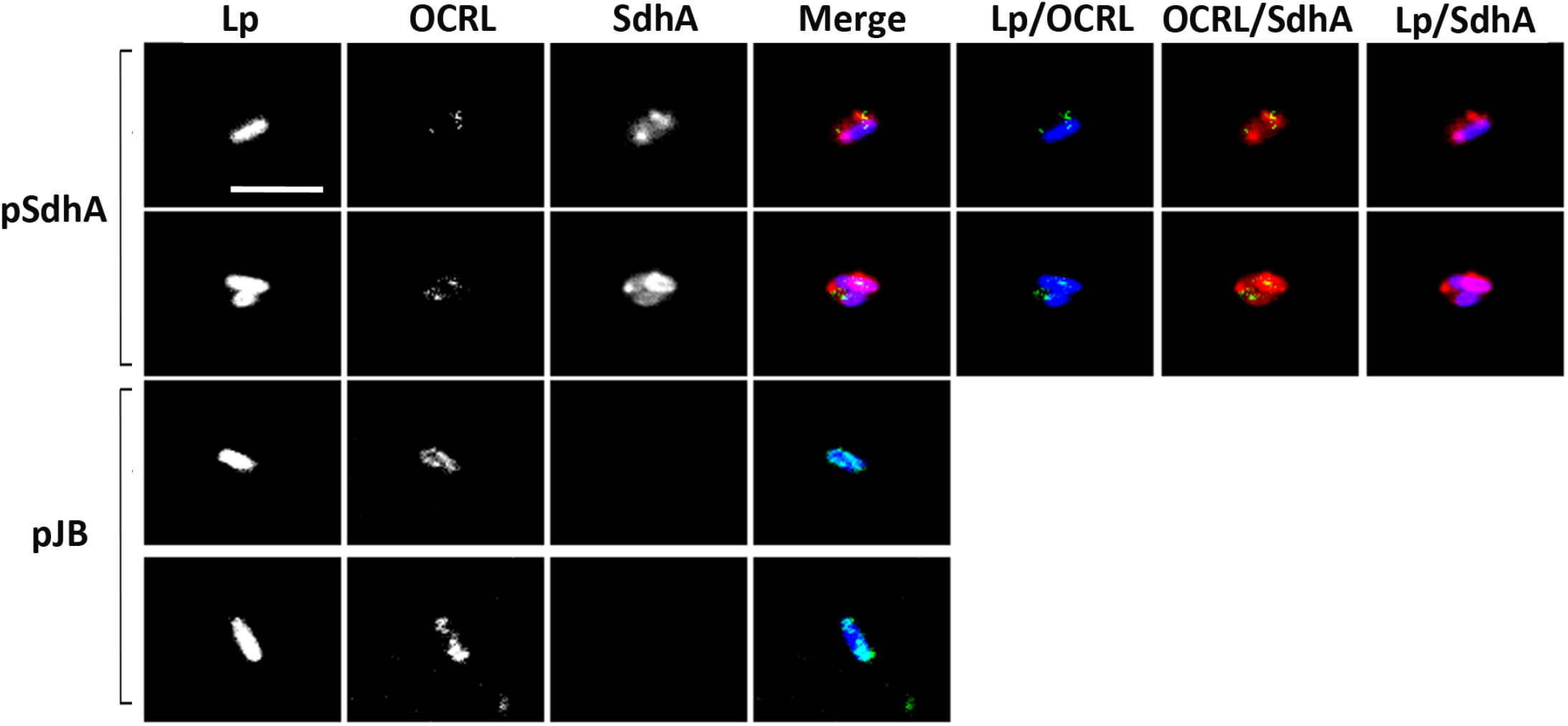
OCRL localization about the LCV surface. Linked to Fig. 6D. Confocal images of vacuoles isolated from infected U937 cells (MOI =10, 3h) with Δ*sdhA* mutant harboring pSdhA or pJB vector. The presence of OCRL and SdhA on LCVs was assessed by immunofluorescence using antibodies against OCRL, SdhA and *L. pneumophila*. (Scale bar, 4 μm)

## STAR METHODS

### KEY RESOURCES TABLE

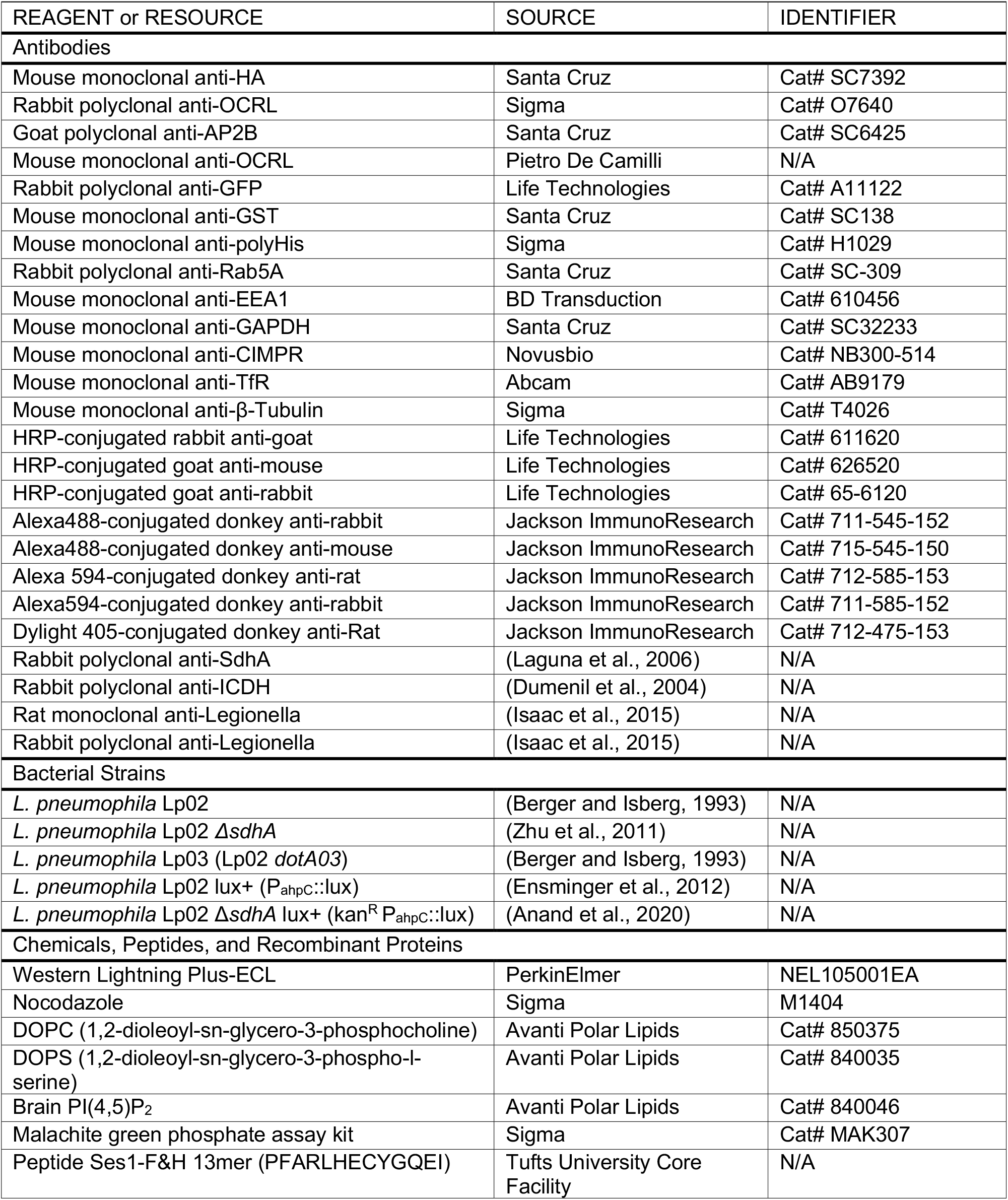
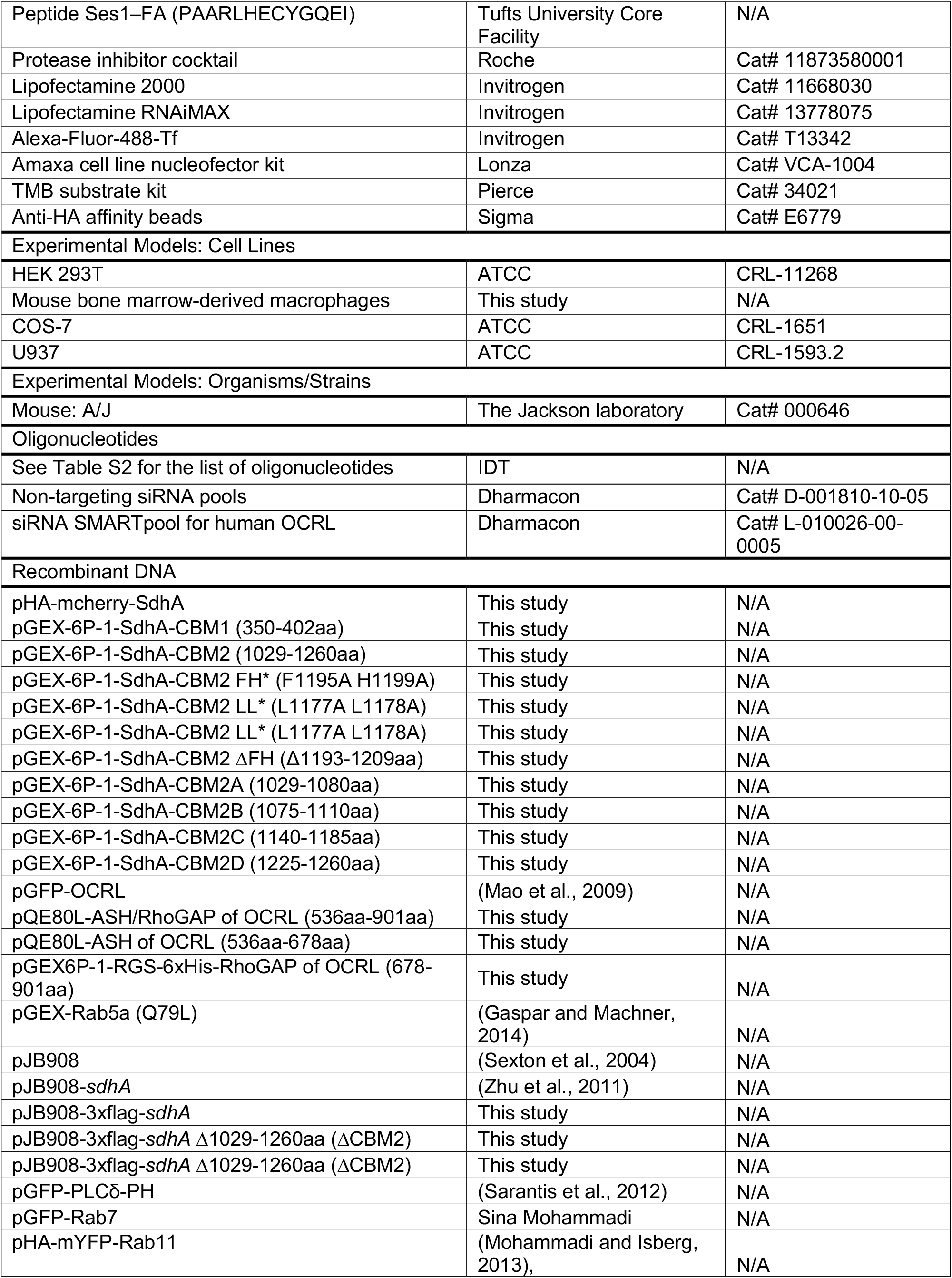
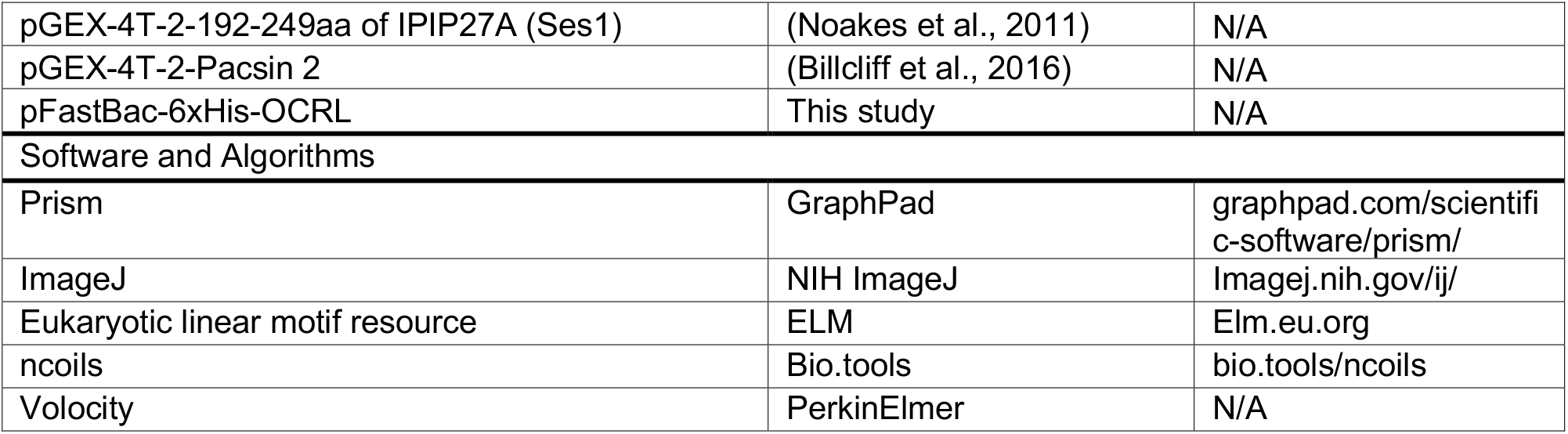

## RESOURCE AVAILABILITY

### Lead Contact

Further information and requests for resources and reagents should be directed to the Lead Contact, Ralph R. Isberg.

### Materials Availability

The materials generated in this study are available upon request.

### Data and Code Availability

The published article includes all data generated or analyzed during this study.

## EXPERIMENTAL MODEL AND SUBJECT DETAILS

### Cell culture

Bone marrow-derived macrophages (BMDMs) from A/J mice were isolated and cultured as previously described (Swanson and Isberg, 1995). COS-7 cells and HEK 293T cells were cultured in Dulbecco’s Modified Eagle Medium (DMEM) (Gibco) supplemented with 10% heat-inactivated FBS (Gibco). U937 cells were maintained and differentiated as described previously (Losick et al., 2010).

## METHOD DETAILS

### Cloning and mutagenesis

The primers used for this work are listed in Table S2. His-Tagged-full length SdhA was constructed in pQE80L while appropriate truncations (CBM1, CBM2, CBM2A, CBM2B, CBM2C, and CBM2D) were generated in pGEX-6P-1. Mutations were generated by PCR using Quikchange (Stratagene). SdhA deletions were generated by inverse PCR (Ochman et al., 1988). For insect cell expression of human OCRL, the full-length cDNA with an amino-terminal 6xHis tag was inserted into pFastBac vector. His-tagged ASH/RhoGAP and ASH domain of human OCRL cDNA were inserted into pQE80L. The amino-terminal-6xHis fused RhoGAP domain of OCRL was inserted into pGEX-6P-1 to generate GST-6His (internal tag) for purposes of improving solubility of the recombinant protein. All constructs were verified by DNA sequencing (Genewiz).

### Co-immunoprecipitation

HEK 293T cells were co-transfected with GFP-OCRL and either HA-mcherry or HA-mcherry-SdhA using Lipofectamine 2000 for 2 days. Lysates were made on ice for 1hr (50 mM Tris-HCl, pH 7.4, 150 mM NaCl, 2% Octylglucoside, protease inhibitor cocktail) and clarified by centrifugation at 16,000g for 20min at 4°C. Immune complexes from the supernatants were adsorbed on anti-HA affinity beads for 2 hr at 4°C. After washing (50 mM Tris-HCl, pH 7.4, 40 mM NaCl, 0.2% Octylglucoside, protease inhibitor cocktail), proteins were fractionated by SDS-PAGE and transferred to polyvinylidene difluoride membranes (Millipore), and probed with the indicated antibodies.

### Pulldown experiments

Pulldown experiments with purified GST or GST-SdhA truncations were performed with the lysates of HEK cells transfected with GFP or GFP-OCRL after 24 hr transfection. The lysates were prepared from a 10 cm dish in 1ml of pull down/lysis buffer (25 mM Hepes–KOH (pH 7.2), 125 mM potassium acetate, 2.5 mM magnesium acetate, 0.4% Triton X-100, and protease inhibitor cocktail), followed by incubation for 3 hr at 4°C with 250 μg of GST-fusion protein coupled to glutathione-agarose (Thermo Scientific). Beads were then washed four times (pull down buffer containing 0.1% Triton X-100) and resuspended with SDS-PAGE sample buffer followed by SDS-PAGE and Western blotting.

### Protein preparations

6xHis-OCRL was prepared from Sf9 insect cells using a baculovirus expression system (Invitrogen) according to the manufacturer’s specification. Cells were lysed (20mM Tris, pH 8, 150mM NaCl, 5mM MgCl_2_, 5mM β-mercaptoethanol, 1% NP40, 10% glycerol, and protease inhibitor) and purified by nickel affinity chromatography (GE healthcare Lifesciences). *E. coli* BL21 (DE3) was used for bacterially-expressed protein, inducing overnight with 0.1 mM IPTG at 18°C. GST fusion proteins were purified on glutathione superflow agarose according to the manufacturer’s protocol (Thermo Scientific). The GST tag of N-terminal GST-6xHis-RhoGAP domain was removed by addition of PreScission protease (GE healthcare Lifesciences) directly to the glutathione beads followed by incubation overnight at 4°C to obtain His-tagged RhoGAP. His-tagged proteins were purified by nickel affinity chromatography. The proteins were then concentrated by ultrafiltration using Amicon filters (EMD Millipore).

### Solid phase binding assay

96-well ELISA plates (Thermo Fisher) were coated with 1ug of recombinant proteins (20 mM NaHCO_3_/Na_2_CO_3_, pH 9.4), washed four times (Tris (pH=7.4)-buffered saline (TBS), 0.1% Tween 20 (TBST)), and blocked in TBS + 5% BSA. Wells were then probed with GST-or His-tagged proteins diluted in TBST containing 0.5% BSA for 1 hr. Wells were washed again with TBST and incubated with appropriate primary antibodies and HRP-conjugated secondary antibodies. Protein interaction was detected by incubating with TMB (Thermo Fisher) as a chromogenic substrate. The reactions were stopped (1M HCl and 5M acetic acid in water) and the resulting absorbance was measured at 405 nm in a microtiter plate spectrophotometer. EC_50_ was calculated using the GraphPad Prism 8 software for windows applying the nonlinear curve fit module.

### Bacterial challenge of mammalian cells

The analysis of intracellular replication at a single cell level and analysis of cytosolic bacteria in BMDMs were performed as previously described (Creasey and Isberg, 2012). To quantify SdhA localization on LCV, SdhA and *L. pneumophila* were probed with anti-SdhA and anti-*Legionella*, respectively, incubated with fluorescent secondary antibodies, and 100 vacuoles were counted. Isolation of vacuoles from *L. pneumophila*-infected U937 was performed as described previously (Vogel et al., 1998). For intracellular growth in COS-7 cells, 2×10^4 cells were seeded in 96-well plates, challenged with *L. pneumophila lux* (MOI 20) for 1 hr, washed three times with Dulbecco’s modified Eagle’s medium (no phenol red) supplemented with 10% FBS, and luciferase production was measured at 48 hr post infection in a microtiter luminometer.

### Imaging

Fluorescence microscopy was performed following standard procedures (Losick and Isberg, 2006) and all antibodies were used according to manufacturer’s procedures. Nuclei were stained using Hoechst stain (Invitrogen). Cells were transfected using Lipofectamine 2000 (Invitrogen) for 24 hr according to the manufacturer’s instructions and fixed with 4% formaldehyde in PBS. For nocodazole treatment, 0.1 µg/ml of nocodazole (Sigma) was added to cells for 20 hrs. Cells were imaged by either Zeiss observer Z1 or Leica Falcon SP8 microscopy. Images were processed using ImageJ or Volocity software (Improvision). The fluorescence intensity of OCRL covering a single LCV and transferrin in the cytoplasm was quantified using ImageJ software after background correction.

### Transferrin uptake and recycling

COS-7 cells-transfected with SdhA or with control vector were incubated in serum-free medium for 1 hr and then exposed to 50 μg/ml Alexa-Fluor-488-Tf (Invitrogen) on ice for 30 min. The cells were transferred to 37 °C and incubated for the times indicated. External Tf was removed by washing with PBS and bound Tf was removed by an acid wash (150 mM NaCl, and 10 mM acetic acid, pH 3.5) followed by washes with PBS. The fluorescence intensity of internalized Tf was quantified by image capturing using a Zeiss Observer Z1 microscope (63x oil objective) and analysis using Image J. To measure recycling, cells were incubated first in serum-free medium and subsequently in medium containing fluorescent Tf for 1 h at 37 °C to saturate the receptor population. After extensive washing with HEPES-buffered DMEM, the recycling of Tf was followed by incubating the cells in the presence of complete medium for 40 and 60 min, at 37 °C. The cells were then acid washed before fixing.

### Lipid phosphatase assay

To measure phosphatase activity, lipid vesicles containing PI(4,5)P_2_ were generated by extrusion with polycarbonate membranes with pore size of 200-nm diameter as described in Billcliff *et al*. (2016). Lipids DOPC (1,2-dioleoyl-sn-glycero-3-phosphocholine), DOPS (1,2-dioleoyl-sn-glycero-3-phospho-l-serine), and natural PI(4,5)P_2_ were purchased from Avanti Polar Lipids. His-tagged OCRL (50 nM) was incubated with indicated proteins on ice for 20 min in reaction buffer (50mM Tris-HCl, pH7.4, 5mM MgCl_2_). Ses1 C-terminal fragment (192-249) and GST-Pacsin2 were added at 20-fold molar excess of OCRL. The phosphatase reaction was started by addition of 200 μM of PI(4,5)P_2_-containing liposomes. After incubation at 37°C for 20 min, the reaction was stopped by the addition of malachite green solution (Sigma-Aldrich) and the resulting absorbance was read at 620nm.

### RNA interference

COS-7 cells were transfected using Lipofectamine RNAiMAX (ThermoFisher Scientific) according to the manufacturer’s instructions. The siRNA SMARTpool for OCRL and non-targeting siRNAs were purchased from Dharmacon. For OCRL depletion in U937 cells, siRNA was transfected with Amaxa cell line nucleofector kit and Nucleofector II (Lonza) according to the manufacturer’s protocol.

